# Fibrilpaint20 directs the ubiquitin–proteasome system to Tau amyloid fibrils

**DOI:** 10.1101/2024.08.27.609886

**Authors:** Françoise A. Dekker, Júlia Aragonès Pedrola, Emile van Weert, Anne den Breejen, Laura Peters, Adriana Poza-Rodriguez, Guy Mayer, Shalini Singh, Alfred C.O. Vertegaal, Assaf Friedler, Stefan G. D. Rüdiger

## Abstract

Failure of the Protein Quality Control (PQC) System to control the Tau protein leads to amyloid fibrils that are at the basis of various Tauopathies, including Alzheimer’s Disease (AD). Reinstating the PQC for Tau is desirable. Here we show that the PROTAC peptide FibrilPaint20 (FP20) recruits the ubiquitination system to Tau amyloid fibrils. We reconstituted the ubiquitin-proteasome cascade and monitored the stepwise ubiquitin transfer using the microfluidics technique FIDA. To direct ubiquitination specifically to amyloids, we discovered FP20, a bifunctional peptide that connects amyloid fibrils to the E3 ligase CHIP. FP20 mediated ubiquitination of recombinant and patient derived fibrils from various tauopathies, ultimately enabling the 26S proteasome to target CHIP-ubiquitinated Tau fibrils. This shows that FP20 cooperates with the PQC system and can initiate fibril cleavage. This offers a strategy to target amyloid diseases by cooperation with the PQC system exploiting key cellular systems.

## 1. Introduction

The Protein Quality Control (PQC) system controls aggregation-prone proteins in the cell. This system consists of a network of molecular chaperones and degradation machineries tasked with maintaining cellular protein homeostasis ^1–3^. When the PQC system fails, misfolded proteins accumulate and assemble into amyloid fibrils ^4,5^. Aggregation of the microtubule-binding protein, Tau, into intracellular amyloid fibrils is a hallmark in Alzheimer’s Disease (AD) and other neurodegenerative tauopathies ^6,7^. Ultimately, this leads to cognitive decline and motor dysfunction in affected individuals ^7–9^

In healthy neurons, Tau homeostasis is maintained by the main chaperone systems of the cell, Hsp70 and Hsp90 ^10–14^. They bind to Tau and recruit the E3 ligase CHIP, which subsequently ubiquitinates Tau and ultimately leads to degradation by the 26S proteasome ^13,14^. As long as this cascade functions, Tau cannot form amyloid fibrils. With aging, however, PQC capacity declines, which may facilitate the appearance of amyloids ^3,15^. This potentially creates ever bigger challenges for PQC due to growing amyloid fibrils in disease. This raises the question what it takes to restore PQC capacity for Tau once fibrils have formed. Although Hsp70 can, in principle, disassemble *in vitro*-generated Tau fibrils, this process is incomplete ^16–18^. Thus, the potential of the chaperone machinery to restore cellular protein homeostasis once fibrils have formed is limited.

Tau is an intrinsically disordered protein, which makes the binding sites of the Hsp70 and Hsp90 chaperones easily accessible for the chaperone machinery. These binding sites reside in the aggregation-prone microtubule binding repeat region of Tau ^19^. During aggregation, these sites may become less accessible for the PQC system ^20–23^. Interestingly, inclusions in patient brains are frequently decorated with ubiquitin, suggesting that the PQC machinery does attempt to mark these aggregates for proteasomal clearance ^24–26^. However, fibrils in the brain of AD patients are predominantly monoubiquitinated, which does not suffice for subsequent action.

It is poorly understood which step in the ubiquitination cascade breaks down first in disease, and what defines the limits of the degradation machinery. Downstream recruitment systems may not effectively engage fibrillar substrates, either due to poor recognition or too few ubiquitin-binding sites available ^27^. Another possibility is that early intermediates of the aggregation pathway may inhibit proteasomal degradation ^28–30^. Or, end-stage fibrils might simply be too large or too stable for the proteasome to process or for lysosomes to surround the fibril ^31,32^. To address these questions, it is key to understand whether ubiquitination can in principle target fibrillar substrates and to which extent it may still render fibrils accessible to proteasomal processing.

Addressing these questions requires tools that allow direct connection of the ubiquitination machinery to amyloid fibrils. Recently, we discovered FibrilPaint1 (FP1), a fluorescently-labelled Proteolysis Targeting Chimera (PROTAC) peptide that specifically recognises fibrils ^33–37^. This work exploits the FibrilPaint concept to recruit the ubiquitin– proteasome system (UPS) to Tau fibrils.

Here, we introduce the PROTAC FibrilPaint20 (FP20), a non-fluorescent derivative of FP1 that specifically binds amyloid fibrils and recruits CHIP via a C-terminal EEVD motif. FP20 recruited recombinant and patient-derived Tau fibrils to the reconstituted E1-E2-E3 ubiquitin cascade. FIDA technology allowed to follow the stepwise travelling of fluorescently labelled ubiquitin throughout this cascade. The 26S proteasome subsequently recognised and also cleaved growing, recombinant TauRD fibrils. Our findings suggest that amyloid fibrils remain accessible to the UPS, which renders the combination of FP20 and CHIP interesting for initiation-controlled removal of pre-formed Tau fibrils.

## 2. Results

### 2.1 FP20 binds to Tau amyloids and CHIP simultaneously

We aim to recruit the UPS system to Tau amyloid fibrils, using the specific recognition of Tau fibrils by FibrilPaint peptides, which is established ^33^. FP1 also carries a C-terminal EEVD motif, which serves as a recognition sequence for recruitment of the E3 ligase CHIP **(Table S1)**. We addressed whether FibrilPaints can engage both CHIP and amyloid fibrils simultaneously. We developed two variants of the FibrilPaint family: FP20 (lacking the N-terminal fluorescein) and FP6 (lacking the C-terminal GSGSEEVD motif) **(Table S1, Fig. 1A**).

**Figure 1.**
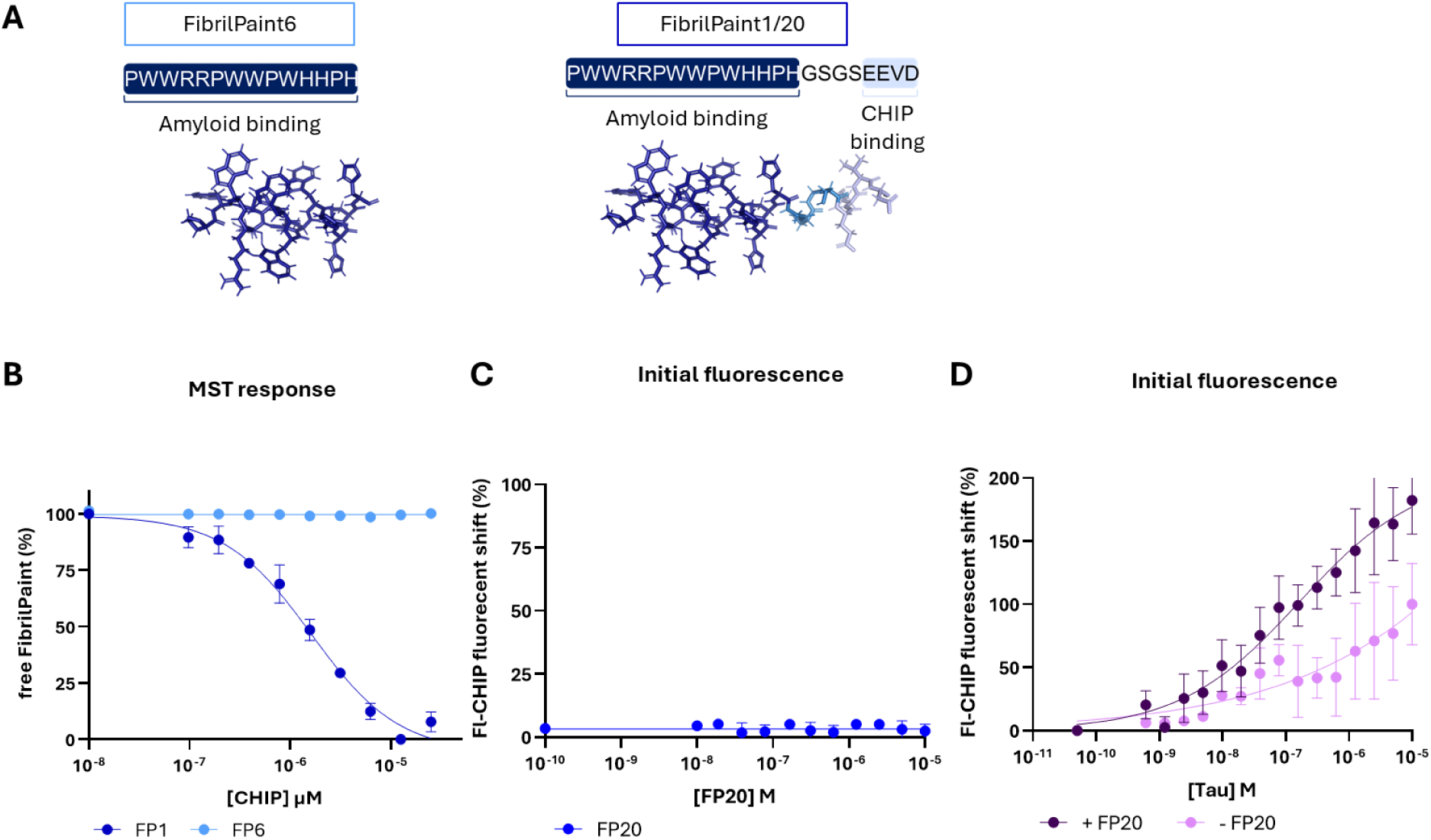
FP1 and FP20 promote binding of the E3 ligase CHIP to amyloid fibrils. (**A**) Sequences and structural representations of FP1, FP20, and FP6. FPs are shown as stick models (blue). Residues 1–14, critical for amyloid binding, are highlighted in dark blue; residues 19–22, mediating CHIP binding, are shown in light blue. (**B**) Binding of CHIP to FP1 (green, N=4) detected by microscale thermophoresis (MST), showing a dose–response curve with an apparent affinity of 1.5 µM. FP6 (blue, N=3) did not elicit a response. (**C**) No initial fluorescence shift was observed when up to 10 µM CHIP bound to FP20. (**D**) Initial fluorescence shift of Fl-CHIP upon binding to TauRD amyloid fibrils. Apparent affinities were above 1.1e-6 M in absence and 1.1e-8 M in presence of FP20.

We titrated CHIP against FP1 and FP6 and measured binding using microscale thermophoresis (MST), monitoring the thermophoretic response. FP1 showed a dose-dependent response to CHIP, with a binding affinity of 1.5 µM (**Fig 1B**). FP6 showed no detectable response up to 25 µM CHIP, demonstrating the requirement of the GSGSEEVD motif for CHIP recruitment by FP (**Fig 1B**).

We investigated whether FP20 can recruit CHIP to Tau fibrils. Fluorescence labelling of CHIP with Atto-565 via an ester reaction (Fl-CHIP) enabled direct detection. The titration of fibrils of the repeat domain of Tau (Tau-RD) into a solution containing Fl-CHIP and FP20 revealed a high affinity of Fl-CHIP to Tau-RD fibrils in the presence of FP20, with a K_D_ of 1.1 × 10⁻⁸ M resulting from MST measurements (Fig. 1D). In the absence of FP20, the K_D_ of Fl-CHIP for Tau fibrils was two orders of magnitude lower (K_D_ 1.1 × 10⁻^6^ M (**Fig 1D**). As FP20 does not alter the fluorescence signal upon binding Fl-CHIP (**Fig 1C**), we conclude that FP20 is an effective recruiter of the E3 ligase CHIP for Tau fibrils.

### 2.1 Flow-based sizing of ubiquitin reveals distinct intermediates of the ubiquitin machinery

Next, we monitored the passage of ubiquitin throughout the degradation cascade. Previously, we pioneered the use the FibrilRuler test, combining FP1 and Flow Induced Dispersion Analysis (FIDA), to monitor fibril length via determining the hydrodynamic radius, R_h_ ^33,38,39^. Here, fluorescein-labelled Ubiquitin (Fl-Ub) allows monitoring the attachment of Ub to various proteins and protein complexes. Ub is a small protein with a predicted R_h_ (R_h_*) of 1.7 nm. If the small Fl-Ub links to another protein, e.g. a Ub-ligase, it results in a larger fluorescent complex, with an increased R_h_ value.

We reconstituted the transfer of the Fl-Ub through the E1-E2-E3 ubiquitin cascade and monitored this via size changes in FIDA experiments ^40,41^ **(Table S2,** Fig. 1A), using UBE1 (E1), UB2K (E2) and CHIP (E3). The structural coordinates of these enzymes enabled the prediction of the expected R_h_ values (with and without ubiquitin), as R_h_ strictly depends on the diffusion properties of the particle, which in turn depends on its dimensions **(Table S2)**. All components of the cascade differ in size, which allowed monitoring of the handover from one step to the next throughout the cascade in order of addition experiments (R_h_ values of UBE1, 4.4 nm; UB2K, 2.3 nm and CHIP 5.1 nm).

The enzymes (500 nM) were sequentially added, while increasing the concentration of the enzyme under investigation to 5 µM, to 10 nM Fl-Ub (Fig. 1B). Free Fl-Ub showed an R_h_ of 1.8 nm, in close agreement with the predicted R_h_* of 1.7 nm **(Table S2)**. The Ub-activating enzyme UBE1 (E1) bound Fl-Ub by forming a thioester bond between the glycine residue of Fl-Ub and a cysteine residue on E1, which is a high-energy bond, requiring ATP ^42^. The R_h_ of Fl-Ub increased to 6.3 nm, consistent with the predicted 5.9 nm of the UBE1-Ub complex. Next, Fl-Ub transferred to the catalytic cysteine residue of Ub-conjugating enzyme E2, UB2K, resulting in an R_h_ of 3.4 nm (R_h_* 3.0 nm). Subsequent interaction of UB2K with the E3 ligase CHIP further increased the R_h_ to 4.7 nm in agreement with, the predicted R_h_ value for CHIP of 5.1 nm. The slightly lower value may reflect the dynamic nature of CHIP. Importantly, Ub transfer through the cascade was strictly ATP dependent: in the absence of ATP, the observed R_h_ remained at 1.8 nm throughout the order of addition experiment, indicating that Fl-Ub did not associate with any ligases. Thus, monitoring the R_h_ of Fl-Ub suffices to demonstrate the transfer of Fl-Ub throughout this E1-E2-E3 cascade.

### 2.2 FP20 enhances ubiquitination of synthetic and patient-derived fibrils

Conjugation of Fl-Ub onto TauRD fibrils by the E3 ligase CHIP came next. Fl-Ub transferred through the E1–E2–E3 cascade under the previously established conditions, with an additional step at the end: addition of recombinant TauRD fibrils as ubiquitination substrate and FP20 as bispecific compound provided a non-covalent bridge between CHIP and the TauRD fibrils (**Fig 2A**). After the journey of fluorescently-labelled ubiquitin through the entire machinery and ultimately onto the fibrils, the R_h_ increased sharply from 1.8 nm to 42 nm (Fig. 2C). This was in agreement with the apparent size of the non-ubiquitinated TauRD fibril in the FibrilRuler test (R_h_ = 44 nm) (Fig. 2C) ^33^. These data indicate that the E1–E2–E3 cascade ubiquitinates TauRD fibrils when using FP20 as bispecific degrader.

**Figure 2.**
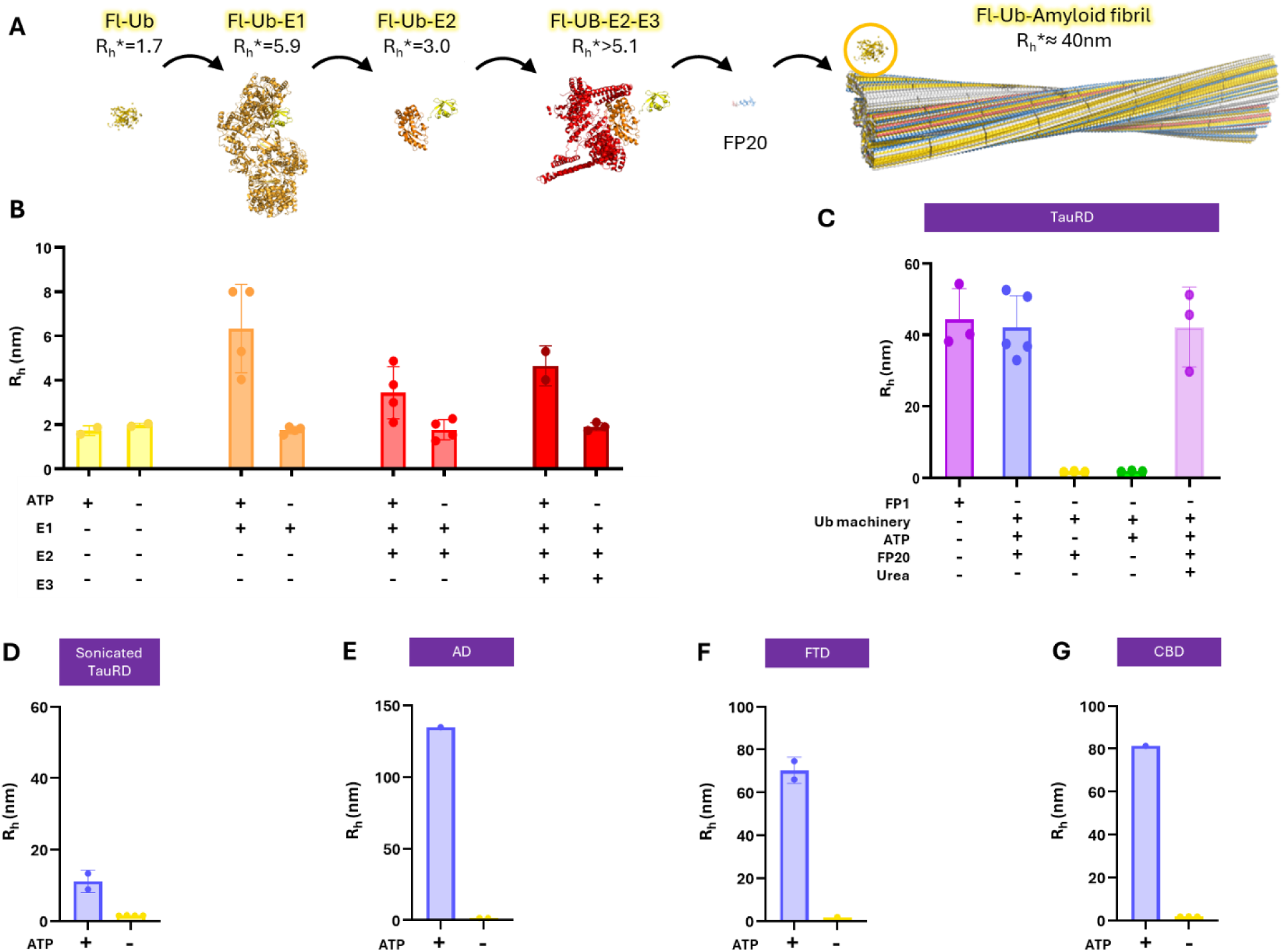
Transfer of ubiquitin over the cascade to TauRD and patient-derived Tau amyloid fibrils. **(A)** Structural representations of the ubiquitin conjugation pathway targeting Tau fibrils, shown to scale. In the presence of ATP, ubiquitin (Ub) is first activated by an E1 enzyme, then transferred to an E2 conjugating enzyme. The Ub–E2 complex is subsequently recognized by an E3 ligase, which binds the substrate for ubiquitin transfer. FP20 facilitates the recruitment of CHIP to the target substrate of Tau amyloid fibrils. For each reaction product, predicted hydrodynamic radius (Rh*) is given, based on available PDB structures. PDB Structures used: Ub, 1UBQ ^43^, E1, 6DC6 ^44^; E2, 2BEP ^45^; E3, 2C2L ^46^. **(B)** Measured hydrodynamic radius (Rh) of fluorescently labelled ubiquitin (Fl-Ub) and components of the ubiquitin cascade, in presence or absence of ATP. Ub alone (yellow); Ub bound to E1 (orange); Ub bound to E2 (red); and the Ub–E2–E3 complex (dark red). Rh increased only in presence of ATP. **(C)** Rh of Fl-Ub measured in the presence of TauRD fibrils, ubiquitination machinery, and FP20. Conditions include TauRD fibrils with FP1 only (purple) ^33^, full machinery with FP20 in the presence of ATP (lilac), full machinery with FP20 in the absence of ATP (yellow), full machinery in absence of FP20 (green) and full machinery with FP20 followed by urea treatment (pink). **(D-F)** Rh of Fl-Ub measured, the full machinery and sonicated TauRD fibrils **(D)**, fibrils purified from patient brains with Alzheimer’s disease **(E)**, frontotemporal dementia **(F)**, and corticobasal degeneration **(G)**, measured in the absence of ATP (yellow) and presence of ATP (lilac).

In absence of ATP, the R_h_ remained 1.8 nm, which corresponds to the size of the free Fl-Ub (Fig. 2C). When repeating the experiment in the presence of all components and TauRD but without FP20, the R_h_ remained 1.8 nm as well, indicating that under these conditions FP20 is essential to facilitate the transfer (Fig. 2C). The complex of TauRD fibrils and Fl-Ub was stable at 4 M urea, the R_h_ value remained at 42 nm (Fig. 2C).This indicates covalent attachment, as urea disrupts most protein–protein interactions while leaving amyloid fibrils intact.

We wondered whether this cascade may also ubiquitinate smaller fibrils. Smaller fibrils appear earlier during the aggregation process, and early targeting of amyloids for degradation may be beneficial for maintaining cellular health ^47,48^. Treatment of TauRD fibrils with a sonicator tip for 30 s resulted in a species with an R_h_ of approximately 10 nm^49^. When adding the E1-E2-E3 cacade, FP20 and ATP, the R_h_ increased slightly to 12 nm, indicating ubiquitination of these smaller fibril species, too (**Fig 2D**).

We next examined whether it could also facilitate ubiquitination of patient-derived fibrils. Patient-derived fibrils differ from recombinant fibrils conformationally ^50^. Tau fibrils are particularly interesting as various Tauopathies are characterised by distinctive fibril morphologies, such as AD, Frontotemporal Dementia (FTD) and Cortical Basal Degeneration (CBD) ^20,23,48^. In AD, tau aggregates into paired helical and straight filaments, in FTD, into narrow filaments; and in CBD, into Type I and II fibrils ^51–53^. These conformational differences alter which residues are surface-exposed or buried, potentially leading to distinct susceptibilities to ubiquitination^23^.

We extracted Fibrils from patient material for AD, FTD and CBD ^33,51–53^. These fibrils underwent the same treatment for ubiquitination as before, followed by monitoring R_h_ of Fl-Ub. For AD, R_h_ extended to 130 nm in presence of ATP (Fig. 2E). For FTD, R_h_ was 68 nm (Fig. 2F) and for CBD 80 nm (Fig. 2G). Thus, ubiquitination was not limited to recombinant structures but could be extended to patient-derived fibrils. This indicates that neither post-translational modifications characteristic of patient-derived Tau fibrils nor the extended fuzzy coat surrounding the fibril core impede FP20-dependent ubiquitination.

### 2.3 Dynamic Light Scattering (DLS) to monitor fibril growth

Having generated ubiquitinated fibrils, the question remained whether these could be processed by the proteasome. Our previous approach, FIDA, relies on detecting fluorescently labelled ubiquitin, but this would leave a single Fl-Ub readout when the proteasome acts on ubiquitinated fibrils. To directly monitor size changes of fibrils in solution, we therefore turned to dynamic light scattering (DLS). DLS is a widely used method to determine particle size in solution, revealing the cumulant radius as readout for the size of particles. DLS can also be used as a relative measure of aggregate dynamics over time ^49,54–57^. We decided to establish the latter approach for Tau aggregation as the presence of large fibrils tends to dominate DLS signal, making absolute cumulant radii unreliable for heterogeneous solutions.

DLS was suitable to monitor TauRD fibril formation over time (Fig. 3). During aggregation, new peaks appeared to the right of the monomer peak, while the relative frequency of the monomer peak decreased (Fig. 3A, B). For each sample, a the DLS measurement revealed a size specific profile revealing a relative cumulant value, termed the apparent R_h_ (R_h__app). Tracking changes in R_h__app indicated whether fibril populations shifted toward more or larger assemblies or smaller species. For example, monomeric Tau showed an R_h__app of 4.2 nm (Fig. 3A), which increased to 74 nm after 24 h of aggregation (Fig. 3B).

**Figure 3.**
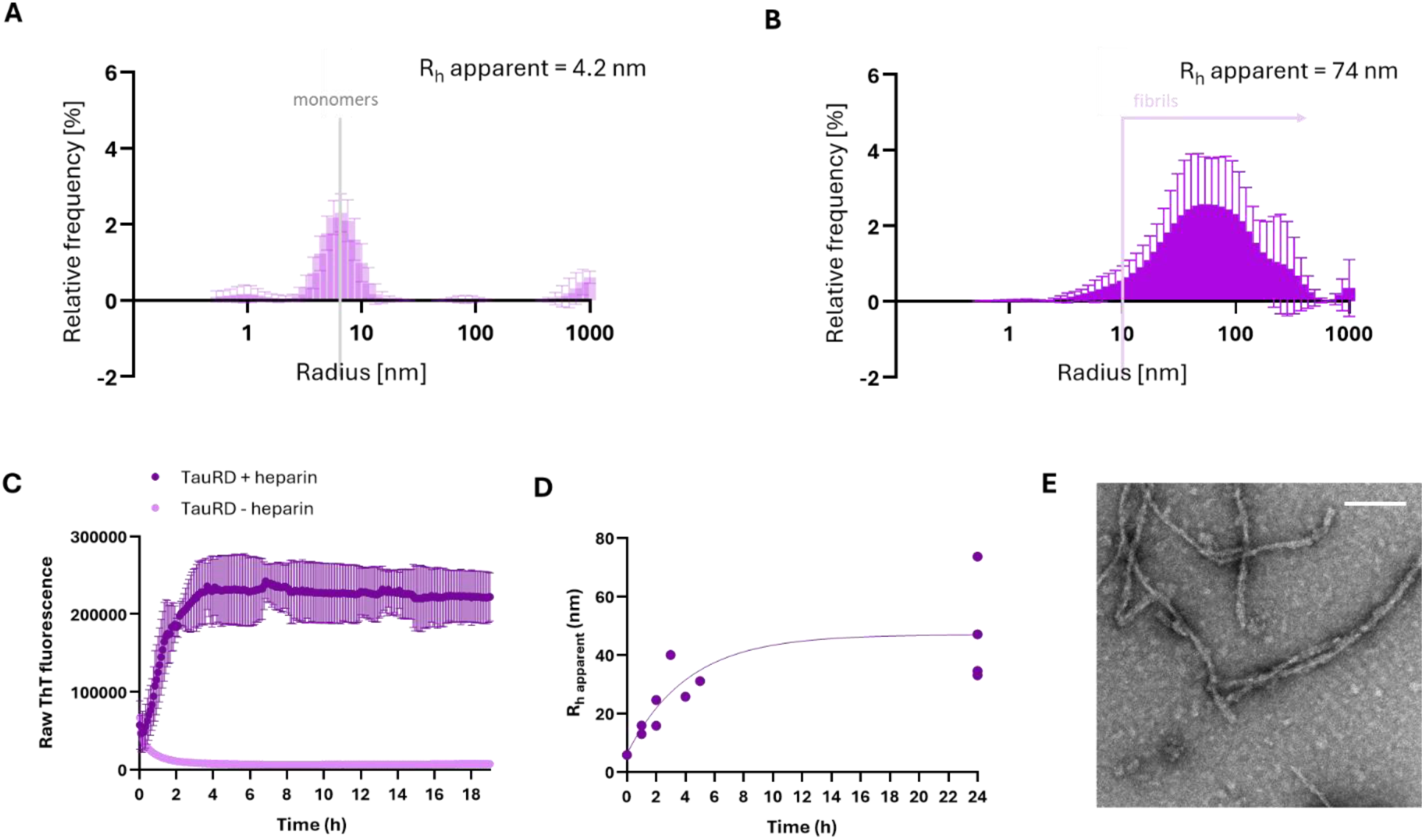
Dynamic light scattering (DLS) to monitor fibril growth. **(A, B)** Representative DLS measurements of TauRD monomers **(A)** and TauRD amyloid fibrils **(B)**. Grey line indicated monomeric species, while the purple line marks the emergence of fibrillar species. Each sample is summarised with the hydrodynamic radius (Rh) as a cumulant value, named Rh_app. **(C)** TauRD aggregation monitored by Thioflavin T (ThT) fluorescence in presence or absence of heparin. **(D)** Corresponding apparent hydrodynamic radii (Rh_app) measured with DLS over time. **(E)** Transmission Electron Microscopy (TEM) image of TauRD fibrils after 24 h of aggregation. White bar in the top right corner reflects a scale of 200 nm.

We compared the R_h__app value obtained by DLS with aggregation monitoring by Thioflavin T (ThT) fluorescence, and Transmission Electron Microscopy (TEM) after 24 h of fibril growth (Fig. 3C**–E**). ThT indicates the presence of the β-sheets formed during aggregation, and the ThT signal rose sharply immediately after the addition of heparin, which triggered the aggregation of TauRD, but not for the condition without heparin (Fig. 3C). ThT fluorescence reached a plateau after 4 h. Like ThT, and R_h__app showed an immediate increase after the addition of heparin (Fig. 3D). Monomers exhibited an R_h__app value of 4.2 nm, which rose to approximately 30 nm after 4 h of aggregation and further increased to 47 nm after 24 h. This was consistent with earlier observations that ThT saturation does not always reflect completion of fibril growth ^33,54,58,59^. TEM analysis of Tau fibrils at the of end of the aggregation process revealed long, singular fibrils, consistent with the DLS results (Fig. 3E). Thus, this DLS approach is suitable to monitor trends in fibril formation.

### 26S proteasome recognises and cleaves ubiquitinated Tau amyloid fibrils

We next tested whether TauRD fibrils can be processed by the 26S proteasome. Polyubiquitination serves as a key degradation signal for both the proteasome and autophagy, with the ubiquitin linkage type determining substrate fate ^60,61^.

The initial setup employed UBE2K (E2-25K), which is suboptimal for our purpose since it specializes in elongating monoubiquitinated substrates ^62,63^. The replacement of this cascade step with the E2 enzyme E2D1 (UbcH5a), a CHIP partner capable of building K48-linked polyubiquitin, allowed to test whether creation of more extensive ubiquitination enables for recognition by a proteolytic clearance system ^64,65^. Increasing the incubation times of the ubiquitination reactions thereby enhanced the likelihood for generating proteasome-recognisable polyubiquitin modifications on amyloid fibrils. We added human 26S proteasomes, freshly purified from HEK293 GP2 cells stably expressing hRpn11-HTBH, subsequently to each experimental condition (0.2 nM) ^66^.

The DLS set-up established for growing TauRD fibrils allowed assessing proteasomal degradation (Fig. 3A, 3B, 4A). To that effect, growing TauRD fibril samples 26S proteasomes incubated for 3.5 h, after which we monitored fibril size via determining R_h__app. Normalising to the R_h__app measured at moment of 26S addition accounted for variability in fibril preparations, such that changes reflect proteasomal activity rather than increased inter-sample heterogeneity (Fig. 4B). Remarkably, the presence of the 26S proteasome lowered R_h__app of Fl-Ub-labelled Tau fibrils over time (Fig. 4C). R_h__app decreased from 90 ± 9 nm to 38 ± 14 nm after 3.5 h. Already after 1 h R_h__app shrank 2-fold, further increasing to a 2.6-fold reduction after 3.5 h. This demonstrates that the 26S proteasome can recognise the ubiquitin-labelled Tau fibrils and cleave them, resulting in fragments that are considerably bigger than the TauRD monomer. Thus, the reaction did not lead to full degradation of the TauRD fibrils. However, our findings indicate that the proteasome can cleave TauRD fibrils after the FP20-dependent labelling in a reconstituted E1-E2-E3 cascade.

**Figure 4.**
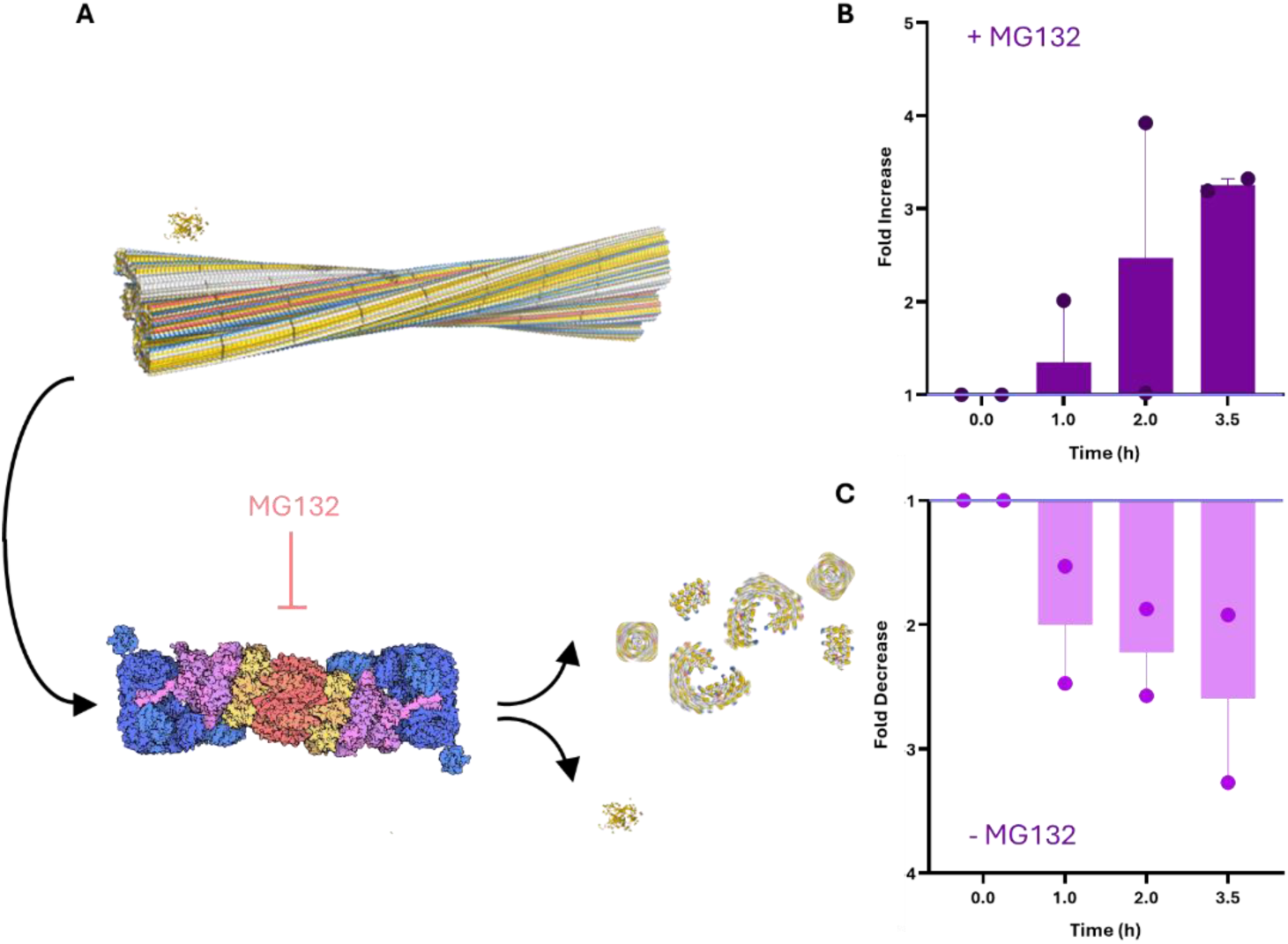
26S proteasomal degradation of amyloid fibrils. **(A)** Schematic representation of ubiquitin-labelled Tau amyloid fibrils recognized by the 26S proteasome, followed by fibril degradation and recycling of ubiquitin. The proteasome is blocked by inhibitor MG132. Structures: ubiquitin (PDB: 1UBQ ^43^); fibril based on Tau PHF (PDB: 5O3L ^51^) and 26S proteasome (PDB: 4CR2 ^67^). **(B, C)** Incubation of TauRD fibrils with 26S proteasome in presence **(B)** or absence **(C)** of MG132. Data were normalized within to reflect the fold change upon addition of the 26S proteasome (t=0). Shown are average results from two independent experiments.

To verify that the degradation activity is specific to the cleavage activity of the 26S proteasome, we did the experiment in parallel in the presence of 50 µM proteasome inhibitor MG132, which we pre-incubated for 10 minutes at 25°C followed by 90 min on ice. In the presence of MG132 and the 26S proteasome, TauRD fibrils showed a time-dependent increase in size, 3.3-fold after 3.5 h (Fig. 4B). Size monitored by determining the R_h__app value increased from 72 ± 13 nm to 236 ± 46 nm. Considering the 2.6-fold decrease in fibril size in the absence of the proteasome inhibitor and the 3.3-fold increase in its presence, the net effect indicates that proteasomal activity reduced the length of TauRD fibrils by approximately 8.6-fold after 3.5 hours. Thus, the specific activity of the 26S proteasome was key to reduce the size of TauRD fibrils.

## 3. Discussion

Here, we showed that the 26S proteasome is capable of cleaving Tau amyloid fibrils, using FP20 as a PROTAC for targeted degradation. By reconstituting the ubiquitination cascade, we showed that FP20 recruits the E3 ligase CHIP to fibrillar Tau, resulting in covalent ubiquitination. This ubiquitination was sufficient to render Tau fibrils recognisable to the proteasome, leading to their partial degradation. These findings establish that mature amyloid fibrils can be engaged by the ubiquitin–proteasome system.

FP20 is a targeted degrader that specifically ubiquitinates Tau amyloid fibrils, both recombinant and patient derived. This distinguishes FP20 from existing Tau targeting strategies, which typically use PET tracer–derived warheads and target total Tau rather than fibrils. Comparable approaches targeting only fibrils are emerging for α-synuclein ^68,69^. Unlike most Tau degraders that recruit von Hippel–Lindau or Cereblon E3 ligases, FP20 employs CHIP as the E3 ligase, expanding the set of PROTAC E3 ligases ^70,71^. CHIP is highly relevant to Tau biology, as it polyubiquitinates Tau in cells ^72^, contributes to Tau clearance ^73,74^, and remains present in the brains of Alzheimer’s disease patients ^75^. The EEVD motif in FP20 offers modularity and can be easily adapted to other targeting strategies. The recognition motif can be readily fused to other protein constructs, enabling the recruitment of additional PQC factors to amyloid substrates.

It is an interesting question what the consequences of proteasomal cleavage of fibrils are for cellular proteostasis. The 26S proteasome shortened fibrils almost by one order or magnitude, but the process was incomplete, suggesting that the proteasome may generate smaller amyloid fragments rather than fully disassembling fibrils. Such fragments may retain or even enhance seeding activity ^76,77^. In the context of disease, 26S proteasomes may be less capable, as the protein quality control capacity declines over time ^3,15,75^ and 26S proteasome function may be impaired ^78,79^, potentially due to inhibition by early aggregate intermediates ^28,30,80,81^.

To understand how FP20 directs ubiquitination, we reconstituted the entire ubiquitin– proteasome system in vitro using purified components. This setup allowed us to trace ubiquitin through the cascade onto amyloid substrates, enabling proteasome-mediated cleavage of fibrils. Cell- or animal based targeted degradation experiments have demonstrated beneficial phenotypic outcomes, such as reduced aggregate load and improved cellular health ^68–71^. However, it remained unclear whether such effects arise directly, through proteasomal engagement with fibrils or inhibition of fibril growth by degradation of monomeric precursors, or indirectly, through alternative routes. By reconstituting the pathway, our work provides mechanistic evidence that mature fibrils themselves can be ubiquitinated and engaged by the proteasome.

Thus, this study shows a proof-of-principle that ubiquitin-proteasome system can act on mature amyloid fibrils, not only their soluble precursors. This demonstrates that even highly structured aggregates remain targetable and can be guided by engineered adaptors, such as FP20. These findings open new opportunities for the development of targeted degradation strategies.

## 4. Materials and Methods

### Expression and purification of TauRD

N-terminally FLAG-tagged (DYKDDDDK) human TauRD (Q244-E372, with pro-aggregation mutation ΔK280) was produced in E. Coli BL21 Rosetta 2 (Novagen), with an additional removable N-terminal His6-Smt-tag (MGHHHHHHGSDSEVNQEAKPEVKPEVKPETHINLKVSDGSSEIFFKIKKTTPLRRLMEAFAKRQGKEMDSLRFLYDGIRIQADQTPEDLDMEDNDIIEAHREQIGG). Expression was induced at OD600 0.8 by addition of 0.15 mM IPTG and incubation at 18 oC overnight. Cells were harvested by centrifugation, resuspended in 25 mM HEPES-KOH pH 8.5, 50 mM KCl, flash frozen in liquid nitrogen, and kept at – 80 °C until further usage. Pellets were thawed at 37 °C, followed by the addition of ½ tablet/50 ml EDTA-free Protease Inhibitor and 5 mM β-mercaptoethanol. Cells were disrupted using an EmulsiFlex-C5 cell disruptor, and lysate was cleared by centrifugation. Supernatant was filtered using a 0.22 μm polypropylene filtered and purified with an ÄKTA purifier chromatography System. Sample was loaded into a POROS 20MC affinity purification column with 50 mM HEPES-KOH pH 8.5, 50 mM KCl, eluted with a linear gradient 0-100%, 5 CV of 0.5 M imidazole. Fractions of interest were collected, concentrated to 2.5 ml using a buffer concentration column (vivaspin, MWCO 10 kDda), and desalted using PD-10 desalting column to HEPES pH 8.5, ½ tablet/50 ml Complete protease inhibitor, 5 mM β-mercaptoethanol. The His6-Smt-tag was removed by treating the sample with Ulp1, 4 °C, shaking, overnight. The next day, sample was loaded into POROS 20HS column with HEPES pH 8.5, eluted with 0-100% linear gradient, 12 CV of 1M KCl. Fractions of interest were collected and loaded into a Superdex 26/60, 200 pg size exclusion column with 25 mM HEPES-KOH pH 7.5, 75 mM NaCl, 75 mM KCl. Fractions of interest were concentrated using a concentrator (vivaspin, MWCO 5 kDa) to desired concentration. Protein concentration was measured using a NanoDrop™ OneC UV/Vis spectrophotometer and purity was assessed by SDS-PAGE. Protein was aliquoted and stored at – 80 °C.

### Expression and purification of CHIP

E3 ligase CHIP was produced with an additional removable N-Terminal His6-Smt-tag ((MGHHHHHHGSDSEVNQEAKPEVKPEVKPETHINLKVSDGSSEIFFKIKKTTPLRRLMEAFAKRQGKEMDSLRFLYDGIRIQADQTPEDLDMEDNDIIEAHREQIGG) in E.coli BL21 Rosetta cells. Expression was induced at OD600 0.8 by addition of 0.2 mM IPTG and incubation at 18 °C overnight. Cells were harvested by centrifugation, resuspended in 25mM Na-Pi pH 7.2, 1500 mM NaCl, flash frozen in liquid nitrogen, and kept at – 80 °C until further usage. Pellets were thawed at 37 °C, followed by the addition of ½ tablet/50 ml EDTA-free Protease Inhibitor and 5 mM β-mercaptoethanol. Cells were disrupted using an EmulsiFlex-C5 cell disruptor, and lysate was cleared by centrifugation. Supernatant was filtered using a 0.22 μm polypropylene filtered and purified with an ÄKTA purifier chromatography System. Sample was loaded into a POROS 20MC affinity purification column with 50 mM Na-Pi pH 8.0, 300 mM NaCl, eluted with a linear gradient 0-100%, 5 CV of 0.5 M imidazole. Fractions of interest were collected, concentrated to 2.5 ml using a buffer concentration column (vivaspin, MWCO 10 kDda), and desalted using PD-10 desalting column to 10 mM HEPES-KOH pH 7.2 buffer., ½ tablet/50 ml Complete protease inhibitor, 5 mM β-mercaptoethanol. The His6-Smt-tag was removed by treating the sample with Ulp1, 4 °C, shaking, overnight. The next day, sample was loaded into POROS 20HQ column with 50 mM HEPES pH 7.2, eluted with 0-100% linear gradient, 12 CV of 1M KCl. Fractions of interest were concentrated using a concentrator (vivaspin, MWCO 5 kDda) to desired concentration. Protein concentration was measured using a NanoDrop™ OneC UV/Vis spectrophotometer and purity was assessed by SDS-PAGE. Protein was aliquoted and stored at – 80 °C.

### Peptide synthesis and purification

The peptides were synthesized using a Liberty Blue Microwave-Assisted Peptide Synthesizer (CEM) with standard Fmoc chemistry and Oxyma/DIC as coupling reagents. The peptide concentrations were measured by UV spectroscopy. The peptides were labelled with 5(6)-carboxyfluorescein at their N’ termini. The peptides were cleaved from the resin with a mixture of 95% (v/v) trifluoroacetic acid (TFA), 2.5% (v/v) triisopropylsilane (TIS), 2.5% (v/v) triple distilled water (TDW) agitating vigorously for 3 hours at room temperature. The volume was decreased by N2 flux and the peptides precipitated by addition of 4 volumes of diethylether at −20 °C. The peptides were sedimented at −20 °C for 30 minutes, then centrifuged and the diethylether discarded. The peptides were washed three times with diethylether and dried by gentle N2 flux. The solid was dissolved in 1:2 volume ratio of acetonitrile I:TDW, frozen in liquid Nitrogen and lyophilized. The peptides were purified on a WATERS HPLC using a reverse-phase C18 preparative column with a gradient IACN/TDW. The identity and purity of the peptides was verified by ESI mass spectrometry and Merck Hitachi analytical HPLC using a reverse-phase C8 analytical column.

### Affinity-based purification of the human 26S proteasome

Protocol was adapted from Wang et al. (2007) Biochemistry. HEK 293 GP2 stably expressing hRpn11-HTBHwere grown in DMEM (ThermoFisher, MA, USA) supplemented with 10% FCS (Serana, Pessin, Germany) and 1% Penicillin/Streptomycin (ThermoFisher, MA, USA) to a confluency of 90% in 7 15-cm cell culture dishes per condition. On the day of lysis, the medium was removed, cells were washed with ice-cold PBS and scraped in ice-cold PBS while the dishes were kept on ice. Cells were lysed in 1 ml lysis buffer (50 mM sodium phosphate pH 7.5, 100 mM NaCl, 10% glycerol, 5 mM MgCl2, 0.5% NP-40. 5 mM ATP, 1 mM DTT, and protease inhibitor tablets (Roche, Basel, Switzerland) were added freshly on the same day of use) per condition and spun at 16.000xg at 4°C for 15 minutes. Supernatant was loaded onto 100 μl High Capacity Neutravidin™ Resin (ThermoFisher, MA, USA) that was washed once with lysis buffer and incubated overnight in a cold room while rotating. After incubation the beads were washed twice with lysis buffer and twice with wash buffer (50 mM Tris-HCl pH 7.5, 10% glycerol. 1 mM ATP and protease inhibitor tablets (Roche, Basel, Switzerland) were added freshly on the same day of use). Proteasomes were cleaved off the beads by incubation with 100 μl elution buffer (Per sample: 100 μl wash buffer containing 5 mM ATP, 1 mM DTT, and 3.2 μM TEV protease. ATP, DTT, and TEV protease were added freshly on the same day of use) at 30°C for 1 hour. Beads were separated from the proteasomes by spinning in Durapore® Ultrafree® – MC – HV 0.45 μm PVDF filters (Merck, MA, USA) at 10.000xg for 3 minutes at 4°C. Aliquots of input, unbound, wash, and elution fractions were mixed with 1X Pierce™ LDS Sample Buffer (ThermoFisher, MA, USA) and 100 mM DTT and loaded on NuPAGE™ Bis-Tris 4-12% gradient gels (ThermoFisher, MA, USA) that were run in 1X MOPS buffer. 26S proteasomes that were used for in vitro experiments were not (flash)-frozen beforehand and always used on the same day of measurement.

### Fibril extraction from human brain

Brain material of FTD and CBD were obtained from the Dutch Brain Bank, project number 1369. Brain material of AD was donated by prof. J. J. M. Hoozemans from the VU Medical Centra Amsterdam.

PHFs and SFs were extracted from grey matter of prefrontal cortex from one patient diagnosed with AD according to established protocols ^51^. Tissue was homogenized using a Polytron(PT 2500E, Kinematica AG) on max speed in 20 % (w/v) A68 buffer, consisting of 20 mM TRIS-HCl pH 7.4, 10 mM EDTA, 1.6 M NaCl, 10% sucrose, 1 tablet/10 ml Pierce protease inhibitor, 1 tablet/10 ml phosphatase inhibitor. The homogenized sample was spun for 20 minutes, at 14000 rpm at 4 °C. Supernatant was collected, and the pellet was homogenized in 10% (w/v) A68 buffer. The homogenized sample was spun once more. The supernatants of both centrifugations were combined and supplied with 10% w/v Sarkosyl, and incubated for 1 hour on a rocker at room temperature. The sample was ultracentrifuged for 1 hour at 100000xg and 4 °C. The supernatant was discarded, and the pellet was incubated overnight at 4 °C in 20 μl/0.2 g starting material of 50 mM TRIS-HCl pH 7.4. The next day, the pellet was diluted up to 1 ml of A68 buffer, and resuspended. To get rid of contamination, the sample was spun for 30 minutes at 14000 rpm, 4 °C. Supernatant was collected and spun once more for 1 hour at 100000xg, 4 °C.

The pellet was resuspended in 30 μl 25 mM HEPES-KOH pH 7.4, 75 mM KCl, 75 mM NaCl, and stored at 4 °C up to a month. Presence of PHF and SF was assessed by TEM.

Narrow filaments were extracted from the grey matter of the middle frontal gyrus from one patient diagnosed with FTD (nhb 2017-019), as described in established protocols ^52^. Fibrils were extracted following the protocol for AD fibrils. After the first ultracentrifugation step, the pellet was resuspended in 250 μl/1g of starting material of 50 mM Tris pH 7.5, 150 mM NaCl, 0.02% amphipol A8-35. The sample was centrifuged for 30 minutes at 3000xg and 4 °C. Pellet was discarded, and the supernatant was ultracentrifuged for 1 hour at 100000xg and 4 °C. The pellet was resuspended in 30 μl of 50 mM TRIS-HCl pH 7.4, 150 mM NaCl, and stored at 4 °C up to a month. Presence of narrow filaments was assessed by TEM.

CBD fibrils were extracted from the grey matter of the superior parietal gyrus of one patient diagnosed with CBD (nhb 2018-007), following established protocols ^53^. Tissue was homogenized using a Polytron(PT 2500E, Kinematica AG) on max speed in 20 % w/v 10 mM TRIS-HCl pH 7.5, 1 mM EGTA, 0.8 M NaCl, 10% sucrose. The homogenate was supplied with 2% w/v of sarkosyl and incubated for 20 minutes at 37 °C. The sample was centrifuged for 10 minutes at 20000xg, and 25 °C. The supernatant was ultracentrifuged for 20 minutes at 100000xg and 25 °C. The pellet was resuspended in 750 μl/1g starting material of 10 mM TRIS-HCl pH 7.5, 1 mM EGTA, 0.8 M NaCl, 10% sucrose, and centrifuged at 9800xg for 20 minutes. The supernatant was ultracentrifuged for 1 hour at 100000xg. The pellet was resuspended in 25 μl/g starting material of 20 mM TRIS-HCl pH 7.4, 100 mM NaCl, and stored at 4 °C up to a month. Presence of CBD fibrils was assessed by TEM.

### TauRD fibril formation

Aggregation of 20 µM TauRD in 25 mM HEPES-KOH pH 7.4, 75 mM KCl, 75 mM NaCl, ½ tablet/50 ml Protease Inhibitor, was induced by the addition of 5 µM heparin low molecular weight. Aggregation reaction was performed using a thermoshaker 37 °C and 600 rpm during 24 hours. For the Thioflavin T (ThT) Assay, reaction was performed in presence of 35 µM ThT and monitored using a ClarioStar Plus at wavelength (440/485 nm) every 5 minutes after 1 minute shaking at 600 rpm.

### Thermophoretic response - Microscale thermophoresis binding assay

Binding between FP1 or FP6 and CHIP was analysed using Monolith (NanoTemper Technologies) Microscale thermophoresis system (MST). Thermophoresis was monitored at a concentration of 50 nM of FP and a dilution series of CHIP, in 25 mM HEPES-KOH pH 7.5, 75 mM KCl, 75 mM NaCl. Samples were transferred to premium capillaries (NanoTemper Technologies) and measurements were performed at 37 °C, with medium blue LED power and at MST infrared laser power of 50% to induce thermophoretic motion. The infrared laser was switched on 1 s after the start of the measurement for a 20 s period. Datapoints used were the MST response at 5 seconds.

### Ligand-induced fluorescence change - Microscale thermophoresis binding assay

CHIP protein was labelled with Alexa Fluor 594 dye through an NHS-esther reaction. Labelling efficiency was determined to be 80 %. Initial fluorescence change was evaluated using Monolith (NanoTemper Technologies) Microscale thermophoresis system (MST) after transfer of the samples to premium capillaries (NanoTemper Technologies). Measurements were performed at 37 °C, with medium blue LED power and at MST infrared laser. First, FP20 was titrated in as a dilution series from 10^-5^ M to a final concentration of 10^-8^M in presence of 20 nM Fl-CHIP. Then, premade amyloids, as described in former sections, were titrated in to 20 nM Fl-CHIP. Maximum fluorescent shift was normalised to 100% as a reference. Next, the same experiments were repeated in presence of 1 µM FP20.

### Ubiquitin Transfer over the ubiquitin cascade

Fluorescein-labeled ubiquitin (Fl-Ub, 10 nM) was sequentially transferred through the ubiquitination cascade (500 nM for E1 (UBE1, Sigma Aldrich), E2 (UBE2K or E2D1, Sigma Aldrich) or E3 (CHIP, purified as described above)) in ubiquitination buffer (40 mM Tris, pH 8.0, 5 mM MgCl₂, 0.05% Tween-20, and 1 mM DTT) in the presence or absence of 5 mM ATP. For each transfer step, the receiving enzyme was added at a 10-fold higher concentration (5 µM) to promote complex formation with Fl-Ub. Reactions were incubated for 1 h at 30 °C. Ubiquitination was assessed on a FIDA1 instrument (480 nm excitation). FIDA1 measurements were performed in capillary dissociation (Capdis) mode and measurement was done at 25 °C. Per run, capillary was cleaned with 1 M NaOH at 3500 mbar for 45 s and equilibrated with buffer (25 mM HEPES, 75 mM KCl, 75mM NaCl, 0.5% pluronic) for 75 s. Sample was injected at 50 mbar for 10 s, and subsequently diffused in buffer at 400 mbar for 300 s.

### Ubiquitination of amyloid fibrils

For ubiquitination of amyloid fibrils, the same ubiquitination reaction was used as described before, together with 500 nM FP20 and the equivalent of 5 μM fibrils. Reactions were carried out in the presence or absence of 5 mM ATP and incubated for 1 h at 30 °C.

FIDA measurements were performed as described before, only the last stap, diffusion in buffer, was done under 75 mbar for 1800 s.

### 26S proteasome cleavage of amyloid fibrils

Ubiquitination reactions were performed as described before. To extend chain formation, UBE2K was replaced with E2D1 (UbcH5a, 500 nM) and reactions were incubated overnight at 30 °C to enhance polyubiquitin chain formation. Human 26S proteasomes were freshly purified from HEK293 GP2 cells and added at 15 µg per reaction in absence or presence of 50 µM MG132. Degradation assays were carried out by incubating ubiquitin-labeled TauRD fibrils with proteasomes for 3.5 h at 30 °C. Proteasomal activity was assessed by dynamic light scattering (DLS) at t=0h, t=1h, t=2h and t=3.5h. To account for variation in fibril preparations, apparent hydrodynamic radii (R_h__app) were normalized to values measured at the time of proteasome addition using GraphPad Prism (v10.4.1).

## Acknowledgements

HEK293 GP2 stably expressing hRpn11-HTBH were kindly provided by the Lan Huang lab, at the UC Irvine School of Medicine. AF thanks The Minerva Center for Bio-Hybrid complex systems and the Saerree K. and Louis P. Fiedler Chair in Chemistry. Measurements on the FIDA1, CLARIOstar® Plus and Monolith were done at the Protein Research Centre of Utrecht University. SGDR was supported by grants of the Campaign Team Huntington and Alzheimer Nederland (No. WE.03-2019-03) and a ZonMW TOP grant (No. 91215084). SGDR and ACOV were supported by the Gravitation Consortium “FLOW” (024.006.036), funded by the Dutch Ministry of Education, Culture, and Science (OCW). FAD and SGDR were supported by NWO TakeOff1 (22073) and BioTech Booster (BIOB25014) grants.

## 6. Author contributions

Conception FAD, JAP, SGDR

Design of the work FAD, JAP, ACOV, SGDR Acquisition FAD, JAP, EW, AB, LP, APR, GM, SS,

Analysis FAD, JAP, EW

Interpretation of data FAD, JAP, SGDR Writing original draft FAD

Revision FAD, SGDR, AF, ACOV

## 7. Supplementary material

**Table S1.**
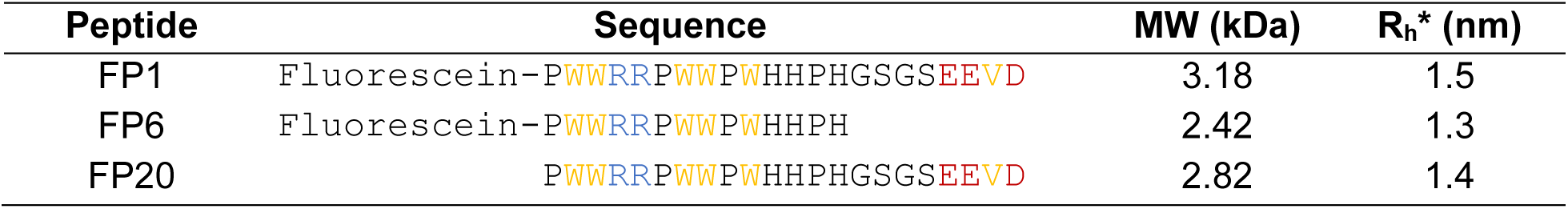
FibrilPaint peptides with corresponding sequences, molecular weight (MW) and predicted Rh (Rh*). Amino acids are coloured as hydrophobic, yellow; positive charge, blue; negative charge, red ^82^.

| Peptide | Sequence | MW (kDa) | $R_h^*$ (nm) |
| --- | --- | --- | --- |
| FP1 | Fluorescein-PWWRRPWWPWHHPHGSGSEEV D | 3.18 | 1.5 |
| FP6 | Fluorescein-PWWRRPWWPWHHPH | 2.42 | 1.3 |
| FP20 | PWWRRPWWPWHHPHGSGSEEV D | 2.82 | 1.4 |

**Table S2:** Used proteins and peptides, used PDB file, predicted hydrodynamic radii (Rh*) and measured hydrodynamic radii (Rh). Predictions are based on the PDB file and BioFIDA software Rh predictor. Unavailable structures indicated with n.a. (not applicable).

| Protein/Peptide | PDB File | Predicted $R_h^*$<br>(nm) | Measured $R_h$<br>(nm) |
| --- | --- | --- | --- |
| FP1 | N/A<sup>33</sup> | 1.5 | 1.7 |
| Ub | 1UBQ<sup>43</sup> | 1.72 | 1.8 |
| E1 (UBE1) | AF-P22314-F1<br>(alphaFold prediction) | 4.37 | n.a. |
| E1-Ub (hUBA1-Ub) | 6DC6<sup>44</sup> | 5.85 | 6.3 |
| E2 (UBE2K/E2-25K) | 2BEP<sup>45</sup> | 2.32 | n.a. |
| E2-Ub (UBE2K-Ub) | 5DFL<sup>83</sup> | 2.98 | 3.4 |
| E2 (E2D1, UbcH5a) | 2C4P<sup>84</sup> | 2.22 | n.a. |
| E2-Ub (E2D1-Ub,<br>UbcH5a- Ub) | 3PTF<sup>85</sup> | 2.52 | n.a. |
| E3 (CHIP) | 2C2L<sup>46</sup> | 5.12 | n.a. |
| E2-E3-Ub | n.a. | n.a. | 4.7 |
| 26S proteasome | 4CR2<sup>67</sup> | n.a. | n.a. |

